# The stress history of soil bacteria under organic farming enhances the growth of wheat seedlings

**DOI:** 10.1101/2023.11.16.567491

**Authors:** Muriel Ornik, Renata Salinas, Giona Antonacci, Martin Schädler, Hamed Azarbad

## Abstract

The effects of stress factors associated with climate change and agricultural management practices on microorganisms are often studied separately, and it remains to be determined how these factors impact the soil microbiome and, subsequently, plant growth characteristics. The aim of this study was to understand how the historical climate and agriculture to which soil microbes have been exposed can influence the growth characteristics of wheat seedlings and their associated bacterial communities. We collected soil from organic and conventional fields with different histories of climate conditions to extract microbes to inoculate wheat seeds. The results showed a positive interaction between conventional farming practices and an ambient climate for faster and higher germination rates. We demonstrate that soil microbial extracts from organic farming with experience of the future climate significantly enhanced above-ground biomass along with bacterial diversity in the seedlings compared to the soil from conventional farming. Our data suggest that soil microbiomes selected by organic farming with a history of future climates can positively impact the aboveground biomass of wheat seedlings and their bacterial assemblages, emphasizing the potential benefits of organic farming practices under changing climate conditions.

## Introduction

The frequency and severity of extreme weather events have been on the rise over the past few decades due to global climate changes and are projected to continue increasing in the future (IPCC 2021). As a result, extreme drought conditions have led to significant reductions in crop yields, ranging from 50% to 70% (Eckstein et al., 2020). This decline in crop production poses a challenge to meet the growing demand for food production, which is expected to increase by at least 70% by 2050 (FAO 2009). The intensive agricultural management, such as conventional farming, involves using high levels of chemicals and fertilizers to increase crop production but can cause land degradation and loss of biodiversity (Foley et al., 2005; Powers and Jetz, 2019). Organic farming methods do not apply synthetic fertilizers or pesticides, which can benefit the overall health of the ecosystem (Bonilla et al., 2012; Lori et al., 2017). Global climate change is likely to strengthen the negative impact of agricultural intensification on different ecosystem functions (Sünnemann et al., 2021). However, we have a limited understanding of how agricultural fields under conventional and organic farming practices can withstand extreme weather conditions and thus ensure food production.

Soil and plant-associated microbes can produce a variety of biological products that can make a significant contribution to crop stress resistance. Climate extremes and agricultural management can both have major impacts on microbes (Azarbad, 2022). The use of high levels of chemicals and mineral fertilizers in conventional farming has been shown to have a significant negative impact on soil characteristics (Dubey et al., 2019), which can adversely affect soil microbiomes (Lupatini et al., 2017). For example, conventional farming has been reported to negatively influence the composition and functioning of the soil microbial community, resulting in a decrease in microbial diversity and biomass (de Vries et al., 2012). On the other hand, agricultural lands that have been managed organically have been shown to have better soil abiotic components, including higher organic matter contents and soil water holding capacity (Lotter et al., 2003), which are essential for microbial growth and function. Accordingly, previous research reported an increase in soil microbial diversity (Hartmann et al., 2015) and abundance (Lori et al., 2017) under this agricultural practice.

In the context of plant microbiomes, beneficial microbes can support the host plants by aiding in nutrient absorption, suppressing pathogens, and protecting against challenging environmental conditions (Li et al., 2019; Azarbad, 2022), thus can directly impact agricultural productivity (Schmidt et al., 2019). By studying 40 different agricultural fields, Ricono *et al*. (2022) reported that organic farming increases the diversity and abundance of beneficial microbes in winter wheat root microbiomes compared to conventional farming. In another example, organic farming has been shown to support a more diverse, complex, and stable microbial network (based on metagenomics sequencing) than that of conventional farming in sugarcane phyllosphere microbiomes (Khoiri et al., 2021). However, as the impact of agricultural practices and the impacts of climate change on soil and plant microorganisms are often studied separately, it is not yet clear how these factors independently or interactively affect plant growth characteristics as well as diversity, and the composition of associated microorganisms.

The aim of this study was to understand how the historical climate and agriculture to which soil microbes have been exposed can influence the growth characteristics of wheat seedlings and their bacterial communities. To achieve that, we collected soil samples from the Global Change Experimental Facility (GCEF). In 2014, the GCEF was established as a field research station located at the Helmholtz-Centre for Environmental Research in Bad Lauchstädt, Saxony-Anhalt, Germany (Schädler et al., 2019). This research station is one of the largest field experiments to evaluate the combined impact of reduced precipitation and warming (as a future climate scenario), as well as different types of land management (such as conventional and organic farming) on various ecosystem processes. This field experiment provides a unique opportunity to study the role of soil microbiome history in shaping the growth characteristics of wheat seedlings. Following soil collections from organic and conventional farming fields (farming history) under ambient and future climate conditions (climate history), we performed an experiment to determine which farming and climate conditions are better in terms of soil microbiomes to improve wheat seedling growth. Specifically, we want to address the following questions: can soil microbiomes selected by a different history of farming and climate affect the growth traits of wheat seedlings? and to what extent does the history of soil microbiomes shape seedlings’ bacterial communities? We hypothesized that the microbiome from inoculum associated with the organic farming soil under historical climate stress contains higher diversity and beneficial microbes that can positively impact the growth characteristics of wheat seedlings. These goals were reached by analyzing important growth traits of wheat seedlings, which are crucial for crop establishment (germination rate and time as well as seedlings above-ground biomass) and bacterial communities from inoculum (soil microbial extracts) and seedlings.

## Material and Methods

### Soil sampling

We took advantage of GCEF, which is a large multi-year field experiment in Saxony-Anhalt, Germany, designed to assess the effects of global changes on various ecosystem processes across a range of land-use types and intensities (Fig. 1). Detailed information about GCEF is available in Schädler et al. (2019). This experimental field includes 50 plots (400 m2 each) arranged in 10 main-plots (5 plots per main-plots). Five different types of land use are established in each main-plot, including (1) conventional farming; (2) organic farming; (3) intensively used grassland that is frequent mowed; (4) extensively used grassland that is moderately mowed; and (5) extensively used grassland that is utilized for moderate sheep grazing. In conventional crop fields, a crop rotation includes winter rape, winter wheat, and winter barley using mineral fertilizers and pesticides (see Schädler et al. 2019 for details). On the organic crop fields, winter rape is replaced by legumes (alternating alfalfa and white clover) as the main fertilization (besides rock phosphate). No pesticides have been applied under organic farming.

**Figure 1.**
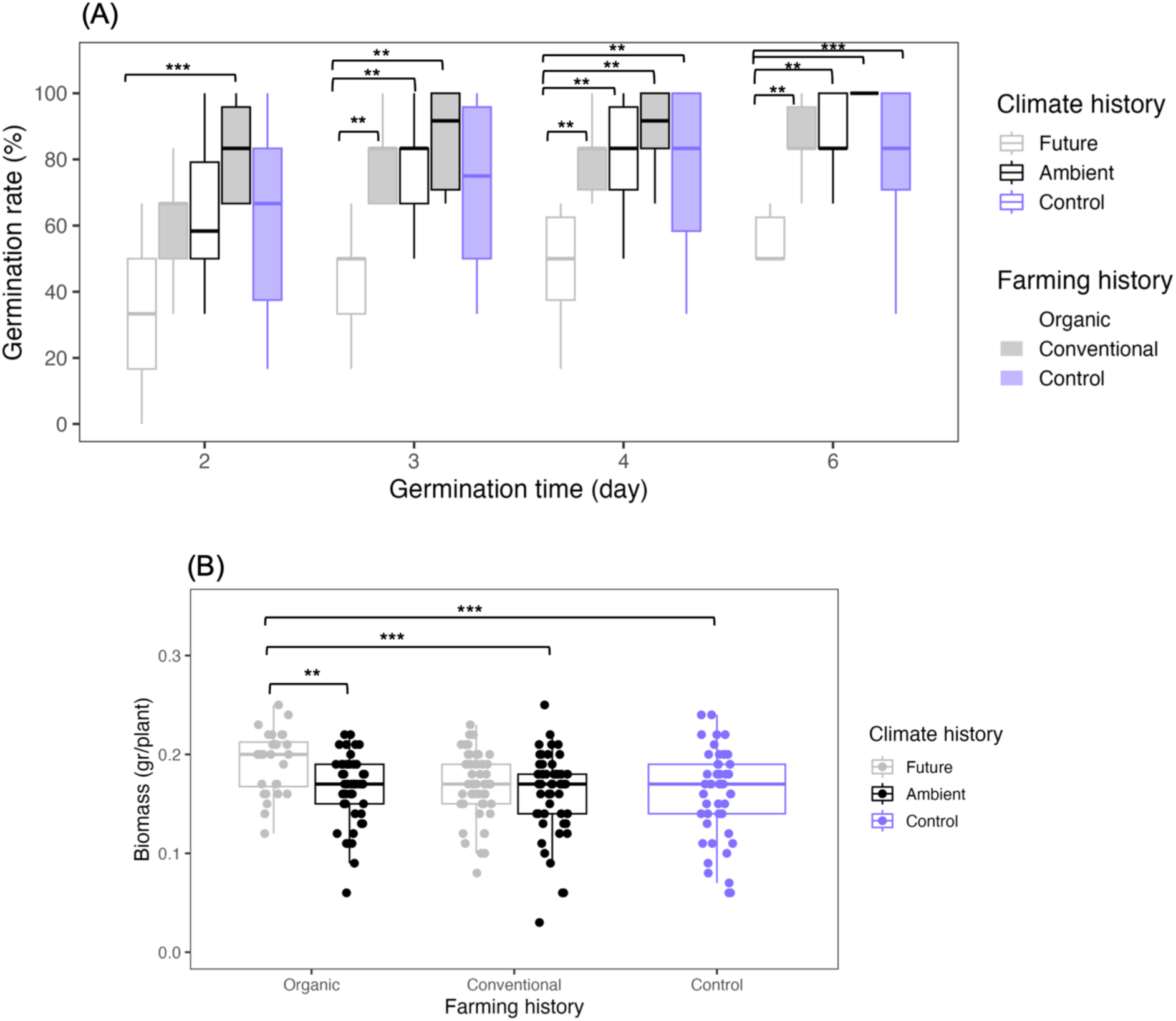
(A) Germination rates and germination times and (B) the above-ground biomass of each seedling where seeds were inoculated with soil microbial extracts from different farming (organic vs. conventional) and climate (future vs. ambient) histories and non-inoculated control plants. Statistical significance was determined by Kruskal—Wallis test (between two levels of farming and climate types and control plants) followed by Dunn’s post-hoc test with Bonferroni method for p-value adjustment. For Germination rates, Kruskal—Wallis test was performed for each germination time separately. Significance code: ‘***’ P ≤ 0.001 ‘**’ P ≤ 0.01 ‘*’ P ≤ 0.05.

Half of the main-plots are subjected to the future climate regime projected as a mean scenario for 2070-2100 in such a way that plots are equipped with mobile roofs (5 m height) and side panels together with an irrigation system. In future climate plots, precipitation is reduced by around 20% in the summer months and increased by around 10% during spring and autumn. Furthermore, the shelters and panels are automatically closed from sunset to sunrise on these plots to increase the temperature in all seasons during the night up to 2.5 °C with a mean increase throughout the year by ∼0.6°C. The rest of the blocks are exposed to the ambient climate. The climate manipulations in the GCEF started in the spring of 2014. Soil samples were collected in August 2021 from future and ambient plots associated with organic and conventional farming systems, representing different soil and microbiome stress histories. For both types of crop fields, the previous crop on the plots was winter wheat (RGT Reformed). Twenty soil sub-samples (2 farming history × 2 climate history × 5 replicates) were collected (0.5-0.6 kg from each replicate) from the topsoil layer 0-15 cm depth), mixed (to have one representative sample), and then transferred to the lab. This resulted in ∼ 3 kg of fresh soil for each soil history type. The selected soils have similar soil types and textures but differ in terms of farming and climate histories. All soil samples were kept moist and cold (at 4°C) till the beginning of the experiment.

### Extraction of the soil microbiome

From each soil history type, microbial extractions were carried out by mixing 5 g of soil (taken randomly from 3 kg of fresh soil) with 45 ml of sterilized PBS buffer in a 250 ml autoclaved Erlenmeyer flask for one hour on the shaker at 150 rpm speed. Next, soil suspensions were transferred into 50 ml Falcon tubes and placed into the centrifuge for 5 mins at 2000 rpm to separate microbial suspension from soil particles. 1 ml from each of these four microbial extractions was immediately taken and stored at −20°C for downstream analysis (e.g., DNA extraction and sequencing assays). This procedure was repeated four times per soil history type and then microbial extractions pooled together before inoculating the seeds. We previously demonstrated that such microbial extraction method led to similar microbial communities in the soil before extraction and microbial suspensions (Giard-Laliberté et al., 2019), which was the case in other studies (Walsh et al., 2021). However, to further confirm whether microbial extraction was successful, we performed the following two steps in parallel. First, 50 μl of microbial suspensions from each soil history were spread on LB and R2A medium agar plates and cultured for 3-5 days at 28°C. We observed complete coverage of the lawn; no colonies were detected in PBS controls. Second, to have an idea about what proportion of bacteria can grow on agar media, we prepared a serial dilution from each soil extract, which allowed us to detect and extract distinguishable bacterial colonies (based on the size, color, and shape of each colony). We then cultured and identified these colonies based on Sanger sequencing of the 16S (LGC Genomics GmbH, Berlin, Germany). These results are shown in Supplementary Figure S1. It is important to point out that, besides microbes, some water-extractable soil nutrients, even though diluted, may have also coextracted from the respective soils with this extraction approach. To check for these factors, soil suspensions were sent to Eurofins Umwelt Nord GmbH (Göttingen, Germany) to measure several important nutrients, including total nitrogen (TNb), Potassium (K), Phosphate (PO4 -3), and total phosphorus (TP).

### Experimental design

Seeds of wheat (*Triticum aestivum* L.) were used in this study. To inoculate the wheat seeds with the soil microbial extracts, microboxes containers (Paungfoo-Lonhienne et al., 2010; Helletsgruber et al., 2017) were sterilized and filled with 200 ml of Murashige Skoog plant agar (1.1 g Murashige Skoog from Duchefa, 5.5 g plant agar per liter distilled water). We did not surface sterilize the seeds before microbial inoculation as we wanted to take into account the possible interactions between the indigenous seed (epiphyte) and inoculum microbiomes. Six seeds were placed in a sterile microbox where each seed was treated with 400 µL of soil microbial extracts by pipetting liquid on the seed and its surrounding area. Control (non-inoculated) seeds received 400 µL of sterile PBS buffer. Therefore, this experiment consists of 5 treatments (four soil microbial extracts plus PBS buffer as control). The sterile microboxes were placed in the growth chamber under controlled conditions of 16:8 h (light: dark cycle), 22-24°C, and 800 µmol m-2s-1 photon flux density. Each treatment was replicated 10 times in a randomized design (5 treatments × 10 replicates = 50 pots). Germination rates and germination times were measured over a period of 6 days. The germination rate was expressed as a percentage where we counted the number of germinated seeds in each treatment and divided it by the total number of planted seeds (6), multiplied by 100. At the end of the experiment (8th day), the plants were carefully removed from the container, the plant agar was discarded, and the seed and roots were cut with a sterile scalpel. The fresh weight of the above-ground biomass (shoot samples) of each seedling was measured for individual plants in each microbox. Following biomass measurements, shoot samples were stored at −20°C prior to DNA extraction.

We decided to grow plants in sterile microboxes instead of soil mesocosms because one of our main aims was to track how fast plant seeds can be germinated with respect to inoculated soil microbiomes. Using microboxes we could monitor the seeds and seedlings throughout the experimental period since day 1. We were aware that only a fraction of soil microbiomes would be able to grow in sterile agar boxes, and thus, only culturable microbes may affect the seedling’s growth parameters. That is why, as mentioned earlier, we also cultured and sequenced those culturable bacteria to have an overview of the proportion of extracted microbes that are able to grow on this media. There are examples where other types of setups (e.g., germination paper) were used to study the effect of the inoculation of wheat seeds with soil microbial extracts on shaping the seedling’s microbiomes (Walsh et al., 2021).

### DNA extraction, amplicon sequencing, and data processing

For extracting DNA from seedlings, 5 replicates (out of 10) were randomly selected within each treatment, where shoot samples of all seedlings in each microbox were ground into a powder using liquid nitrogen with a mortar and pestle. DNA was extracted from 0.1 g of seedlings or 1 ml solution of microbial extract using a phenol-chloroform extraction method (Dellaporta et al., 1983). This resulted in 25 DNA samples from seedlings and 16 samples from soil microbial extracts. Further information on DNA extraction is presented in Azarbad *et al*. (2018). DNA samples were sent to LGC Genomics GmbH (Berlin, Germany) for libraries preparation and Illumina MiSeq (paired-end) sequencing. For the bacterial 16S rRNA gene, the V3-V4 region was amplified using primers 341F (CCTACGGGNGGCWGCAG) and 785R (GACTACHVGGGTATCTAAKCC). Sequence-specific peptide nucleic acid (PNA) clamps, as recommended by Fitzpatrick *et al*. (2018), were used in order to block the amplification of plant-derived DNA and reduce host mitochondrial and chloroplast DNA during amplification. Sequence data were analysed following procedures described by Walsh *et al*. (2021). Raw reads were processed using a DADA2-based bioinformatic pipeline version 1.10.1 (Callahan et al., 2016) in the R software. Briefly, primer sequences were removed, and reads were truncated to 250 and 200 bp for forward and reverse reads (maxEE = c(2,2), maxN = 0, truncQ = 2, rm.phix=TRUE), respectively. Chimeric sequences were identified and removed with the removeBimeraDenovo function of DADA2. For taxonomic affiliations of the resulting amplicon sequence variants (ASV), a naive Bayesian classifier was performed based on the SILVA database v138. ASVs unassigned at the phylum level, together with chloroplast and mitochondrial reads, were removed from the dataset. Before further analysis, the following filtering criteria were applied: samples should have more than 1,000 ASV reads, and any ASVs with less than five reads in a given sample were removed. Furthermore, any ASV that was found in only one sample was discarded from the data set. For the calculation of alpha diversity, the data were normalized on the basis of sequencing depth. For beta diversity, we normalized the data set based on the relative abundance of ASVs in each sample (Walsh et al., 2021). Fastq files are deposited in the NCBI Sequence Read Archive (the BioProject accession PRJNA526458).

### Statistical analyses

Statistical analyses were performed using R software (v 4.4.2, The R Foundation for Statistical Computing). Parametric testing assumptions (the normality and homogeneity of variances) were not met for germination rate and seedling biomass data. Therefore, the Kruskal-Wallis test and post-hoc Dunn’s test with Bonferroni correction (for multiple testing) were used to test the impact of farming and climate histories on seedlings’ growth characteristics. Shannon’s diversity index was calculated to investigate the impact of farming and climate histories on the bacterial diversity of soil microbial extract and seedlings. An analysis of variance (ANOVA) was conducted to determine the potential impact of farming and climate histories on Shannon’s diversity index, considering both direct and interactive effects. We conducted principal coordinate analyses (PCoA) and the Permanova test (through the ‘*adonis*’ function) to evaluate the effect of experimental factors on the bacterial community composition of soil microbial extract and wheat seedlings. Our analysis was based on the relative abundance of ASVs, using Bray-Curtis dissimilarity. For each farming and climate history, to determine if soil microbial extracts are more similar to inoculated seedlings compared to non-inoculated control plants, we calculated the Bray-Curtis dissimilarity for each extract and corresponding plants and tested the significance of the differences using the Wilcoxon ranked test. Additionally, we compared the number of shared and unique ASVs between the bacterial communities of inoculated seedlings and their respective inoculum (soil microbial extracts) to the number of shared and unique ASVs between the communities of non-inoculated control plants and the same inoculum (Giard-Laliberté et al., 2019). We also used ANOVA to analyze the impact of each experimental factor on water-extractable nutrients and the relative abundance of dominant bacteria at phylum and order levels.

## Results

### Water-extractable nutrients in soil suspensions

The total nitrogen concentration (TNb) was higher in soil extract from organic than in conventional farming. Soil extracts from ambient climates contained a higher amount of TNb than in future climates under both farming management (Fig. S2A). The opposite pattern was observed for total phosphorus (TP) and phosphate (PO4 -3) concentrations, where soil extracts from conventional farming had more amount of these nutrients than organic farming. In addition, soil suspension from organic farming under ambient climate appeared to have the lowest concentration of these nutrients (significant farming × climate interaction; Fig. S2C and D). Potassium (K) had, in general, a higher concentration in soil extract originating from future climate, independent of farming systems (Fig. S2B).

### Seedlings growth parameters

The germination results showed that seeds inoculated with soil microorganisms extracted from conventional farming, with a history of ambient climate, exhibited the highest and fastest rate of germination when compared with other treatments (Fig. 1A; Table S1). Conversely, microorganisms derived from organic farming and exposed to future climate recorded the lowest and slower rate of germination in all treatment combinations (Fig. 1A; Table S1). These results suggest that the germination of wheat seeds depends on the type of agricultural soil and climate conditions, where interaction between conventional farming practices and ambient climate can lead to faster and higher germination. As for seedlings’ biomass, we observed that soil microbial extract from organic farming with experience of future climate significantly enhanced seedlings’ above-ground biomass compared to soil microbes from conventional farming and control treatments (Fig. 1B; Table S1).

### The diversity and structure of bacterial communities in soil extracts and seedlings

No significant independent or interactive effects of farming and climate histories were observed for the bacterial diversity in the soil extracts (Fig. 2A; Table 1). However, seeds inoculated with soil microbial extracts from organic farming showed higher bacterial diversity associated with seedlings than other treatments (Fig. 2B; Table 1). Furthermore, the impact of soil microbe’s climate history was only evident in organic farming, where seedlings that were exposed to soil microbes with a history of future climates displayed a higher level of bacterial diversity (significant interactive effect of Farming and Climate; Table 1 and Fig. 2B).

**Figure 2.**
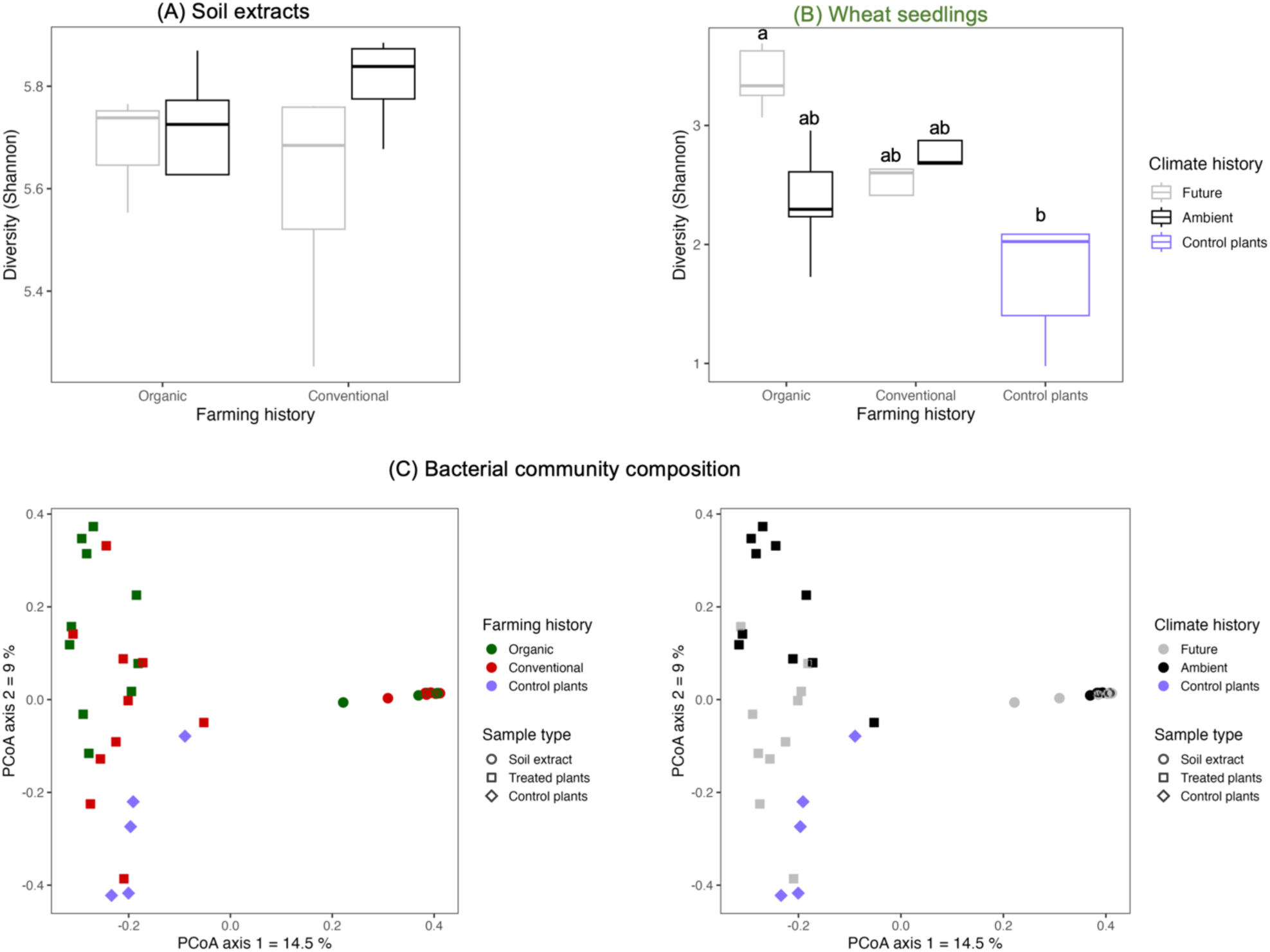
Bacterial Shannon diversity of (A) soil microbial extracts and (B) wheat seedlings. (C) Principal coordinate analyses (PCoA) of Bray–Curtis dissimilarity visualizing the impact of sample type, farming and climate histories on bacterial community composition associated with soil extracts and wheat seedlings.

**Table 1.** ANOVA test for the effects of farming and climate histories and their interactions on bacterial Shannon diversity and bacterial communities of soil extracts and seedlings. Farming refers to the history of soil microbes extracted from organic and conventional agriculture, and climate refers to the history of soil microbes extracted from future and ambient climate under each farming.

|  | Shannon diversity |  |  |  | Community structure |  |  |  |
| --- | --- | --- | --- | --- | --- | --- | --- | --- |
|  | Soil extracts |  | Seedling |  | Soil extracts |  | Seedling |  |
|  | F-value | p-value | F-value | p-value | F-value | p-value | F-value | p-value |
| Farming | 0.062 | 0.808 | 5.305 | **0.014** | 1.826 | 0.087 | 3.421 | **0.001** |
| Climate | 1.381 | 0.265 | 2.206 | 0.153 | 4.672 | **0.001** | 3.986 | **0.001** |
| Farming × Climate | 1.468 | 0.251 | 7.764 | **0.011** | 1.946 | 0.058 | 2.505 | **0.001** |

We conducted a Principal Coordinate Analysis (PCoA) to see how the bacterial community structure was influenced by the experimental factors, considering both the soil microbial extracts and wheat seedlings. These results revealed noticeable differences in the bacterial communities between sample types where soil extract and seedlings clustered separately along the first axis of the PCoA plots (Fig. 2C). The bacterial communities of the seedlings were separated by experimental factors along the second axis of the PCoA plots (Fig. 2C). Due to strong effect of sample types on bacterial communities, we conducted Permanova analyses for soil extracts and seedlings samples separately to confirm PCoA patterns (Table 1). The results of the Permanova analyses indicated that climate history had a significant direct effect on the bacterial community structure of soil microbial extract (Table 1). As for the seedlings, farming and climate histories of soil microbes had significant direct and interactive effects on the restructuring of the seedling bacterial community. Climate history was found to be a major source of variation (higher F value in Table 1) with a clear gradient from ambient to future to non-inculcated control plants along the second axis of the PCoA plots, which was more pronounced under organic farming (Fig. 2C).

### Similarity and shared ASVs between soil extracts and seedlings

We further tested whether there was a greater similarity between soil microbial extracts and inoculated seedlings compared to non-inoculated control plants. The Bray-Curtis dissimilarity was calculated for each extract and its corresponding inoculated plants, as well as for the extract and control plants. Results showed that for organic farming, seedlings that were inoculated with microbes sourced from the soil with past experience of future climate were more similar to each other than control plants (nonsignificant p-value for PC2; Table S2). There were also more shared ASVs between soil extract and seedlings from future climate (16 ASVs; Fig. 3A) than ambient climate. However, for plants introduced to microbes from conventional farming, seedlings showed more similarity (Table S2) and shared ASVs (23 ASVs; Fig. 3B) with their respective soil extract only for the ambient climate. These results indicate that certain members of the bacterial communities were recruited by plants from the soil microbial extract (inoculum) based on the agriculture and climate conditions that soil microbes were exposed to.

**Figure 3.**
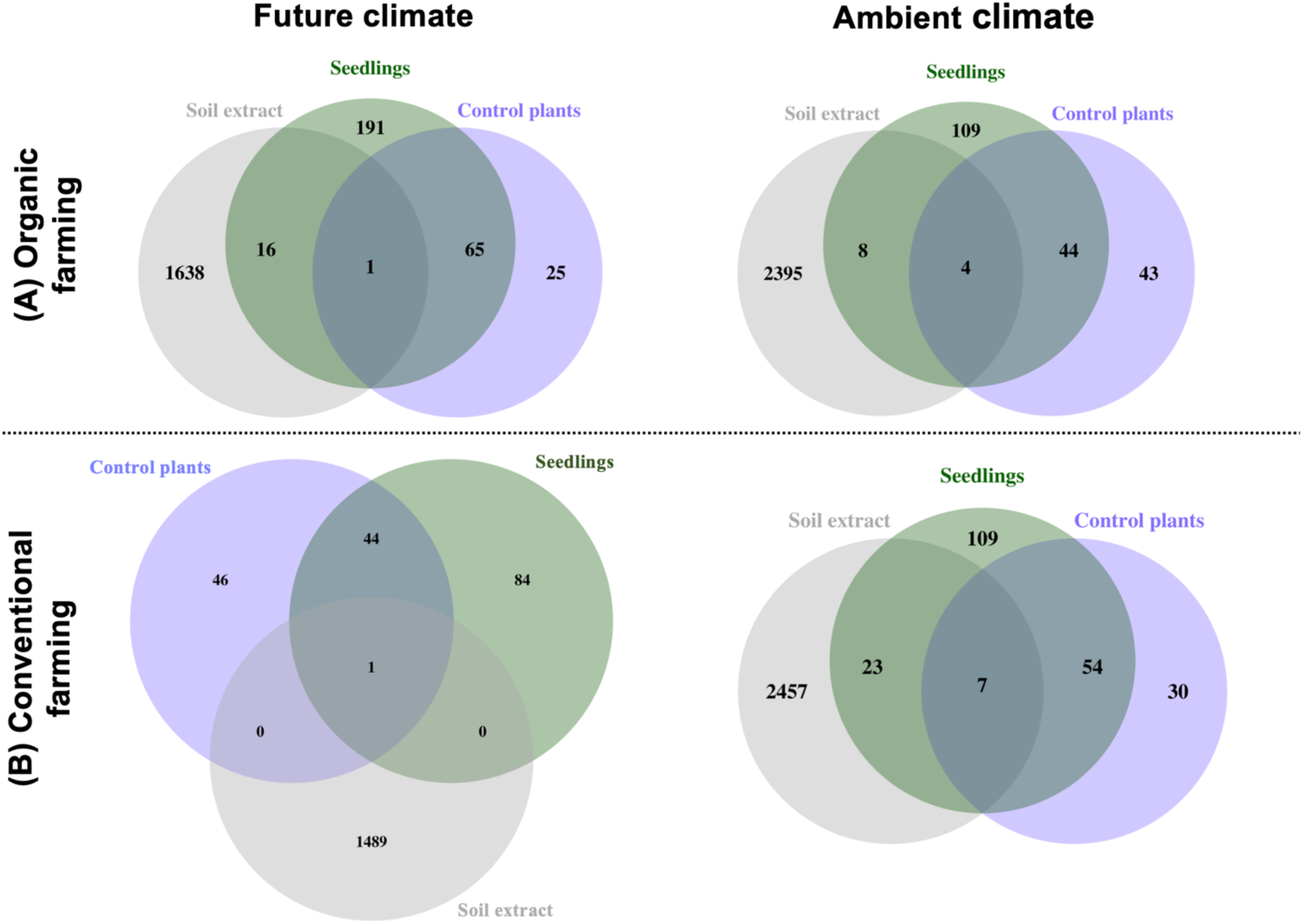
The number of shared and unique bacterial ASVs between the bacterial communities of inoculated seedlings and their respective inoculum (soil extracts) to the number of shared and unique ASVs between the communities of non-inoculated control plants and the same inoculum. The pattern is shown for the plant in which the seed was inoculated with a microbial extract from (A) organic and (B) conventional farming under future and ambient climates separately.

### The composition of bacterial communities

We examined changes in the composition of bacterial communities at the phyla/class (top 10; Fig. S3) and order levels (top 17; Fig. 4) for soil extracts and inoculated and non-inoculated plant samples. At the order levels, in the case of soil extracts, the relative abundance of *Flavobacteriales* and *Xanthomonadales* under conventional farming was significantly higher than for organic farming (Fig. 4; Table S3). Next, we performed ANOVA on soil extracts and treated plants to examine the climate effect for each farming history independently (Table 2). As for soil extracts, the effect of climate was evident for several bacterial orders only for conventional farming (Table 2; Fig. 4). For instance, the relative abundance of *Haliangiales*, *Pseudomonadales*, *Rhizobiales*, and *Sphingobacteriales* were significantly higher in ambient than in future climates (Table 2; Fig. 4). On the other hand, the relative abundance of *Micrococcales* and *Vicinamibacterales* were significantly higher in the future than in ambient climate (Table 2; Fig. 4). These results indicate the effect of climate on bacterial communities, with certain bacterial orders showing significant differences in abundance between ambient and future climates, depending on farming methods. When it comes to treated plants, it was found that the relative abundance of *Burkholderiales* and *Sphingobacteriales* was notably higher in the future compared to the current ambient climate. However, this was only observed in the case of organic farming (Table 2; Fig. 4).

**Figure 4.**
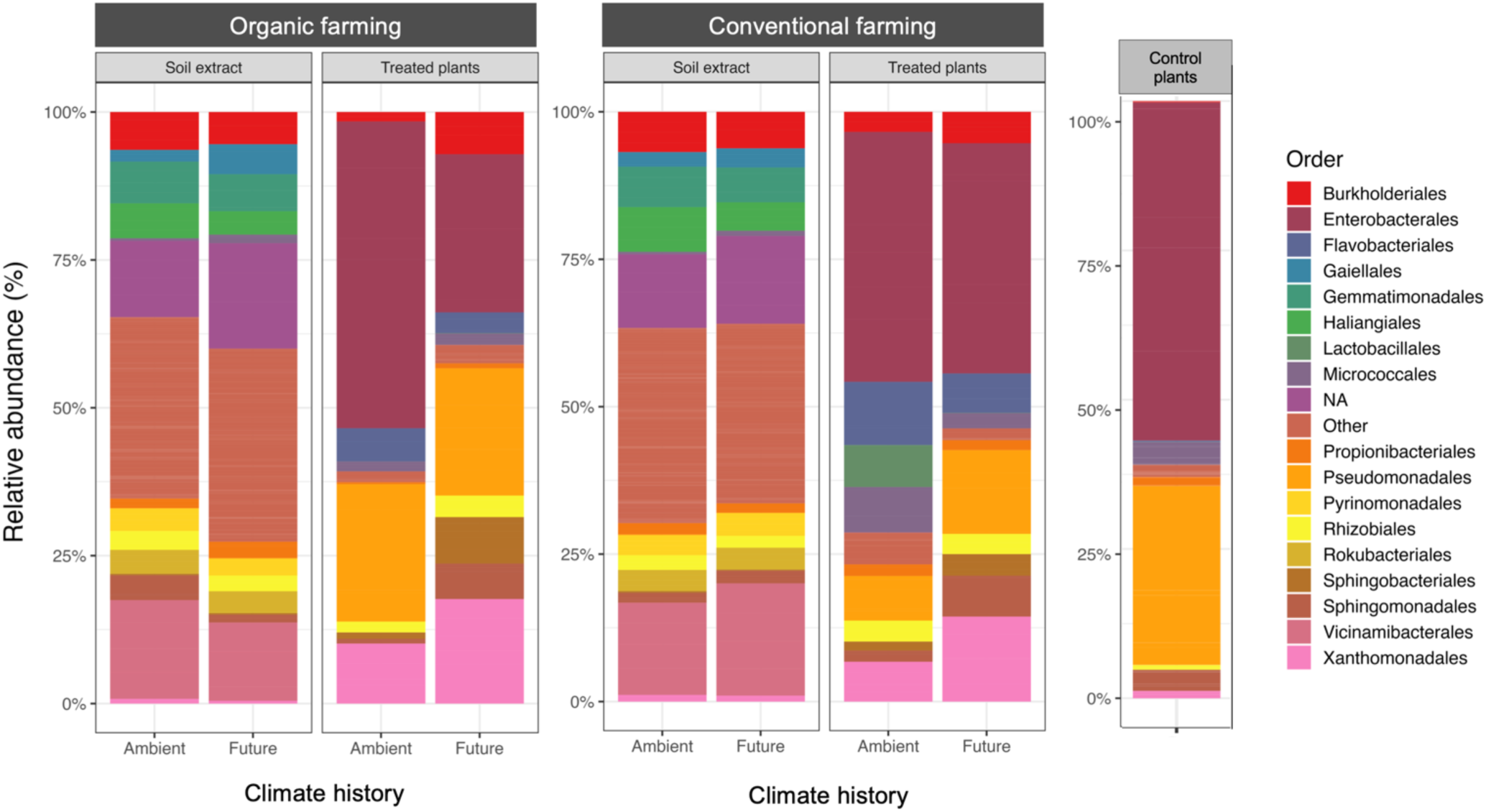
Relative abundance of the most abundant bacterial order associated with soil extracts, non-inoculated (control), and inoculated (treated) plants. The relative abundance is shown according to inoculated plants and their corresponding soil microbial extracts with different farming (organic vs. conventional) and climate (future vs. ambient) histories.

**Table 2.** ANOVA test for the effects of climate histories under organic and conventional farming on most abundant bacterial order associated with soil extracts and seedlings.

|  | Soil extracts |  |  |  | Seedlings |  |  |  |
| --- | --- | --- | --- | --- | --- | --- | --- | --- |
|  | Organic farming |  | Conventional farming |  | Organic farming |  | Conventional farming |  |
| Bacterial Order | F-value | p-value | F-value | p-value | F-value | p-value | F-value | p-value |
| Burkholderiales | 0.345 | 0.582 | 1.802 | 0.228 | 5.12 | **0.053** | 0.944 | 0.36 |
| Enterobacterales | - | - | - | - | 4.101 | 0.077 | 0.033 | 0.861 |
| Flavobacteriales | 0.144 | 0.719 | 0.106 | 0.756 | 0.607 | 0.458 | 0.806 | 0.395 |
| Gaiellales | 1.267 | 0.311 | 1.568 | 0.257 | - | - | - | - |
| Gemmatimonadales | 0.171 | 0.697 | 1.901 | 0.217 | - | - | - | - |
| Haliangiales | 2.052 | 0.211 | 20.273 | **0.004** | - | - | - | - |
| Lactobacillales | - | - | - | - | 3.332 | 0.105 | 1.031 | 0.34 |
| Micrococcales | 1.8 | 0.237 | 9.888 | **0.02** | 0.127 | 0.73 | 1.404 | 0.27 |
| Propionibacteriales | 0.627 | 0.464 | 1.426 | 0.277 | 1.958 | 0.199 | 0.023 | 0.884 |
| Pseudomonadales | 0.303 | 0.606 | 12.479 | **0.012** | 0.023 | 0.882 | 0.873 | 0.377 |
| Pyrinomonadales | 1.696 | 0.25 | 3.089 | 0.129 | - | - | - | - |
| Rhizobiales | 0.105 | 0.759 | 18.08 | **0.005** | 2.088 | 0.186 | 0.005 | 0.947 |
| Rokubacteriales | 0.06 | 0.817 | 0.206 | 0.666 | - | - | - | - |
| Sphingobacteriales | 0.021 | 0.891 | 6.735 | **0.041** | 24.30 | **0.001** | 2.114 | 0.184 |
| Sphingomonadales | 0.685 | 0.446 | 4.167 | 0.087 | 1.649 | 0.235 | 0.709 | 0.424 |
| Vicinamibacterales | 1.484 | 0.278 | 10.275 | **0.018** | - | - | - | - |
| Xanthomonadales | 2.62 | 0.166 | 0.937 | 0.37 | 2.576 | 0.147 | 3.649 | 0.093 |

## Discussion

In this study, we took advantage of a large multi-year field experiment where we collected soil from organic and conventional fields with different climate conditions histories to extract microbes to inoculate wheat seeds. The main aim was to study how agricultural practices and climate conditions independently or interactively impact the growth parameters of wheat seedlings and their associated bacterial communities. We showed that seeds inoculated with soil microbial extracts with a history of conventional farming and an ambient climate exhibited the highest and fastest germination rate. In contrast, seeds inoculated with microorganisms from organic farming and the future climate revealed the lowest and slower germination rate. We further showed that soil microbial extract from organic farming with experience of the future climate significantly enhanced above-ground biomass along with the diversity of bacterial community associated with the seedlings compared to conventional farming. This suggests that specific soil extractable microbes from organic agriculture under future climate positively impact seedling aboveground biomass as well as their associated bacterial communities, emphasizing the potential benefits of organic farming practices under changing climate conditions.

Soil collected from experimental field plots under future climate experienced more changes in precipitation and increased temperature (on average by 0.55°C daily mean temperature and 1.1°C minimum temperatures) in comparison with ambient plots (Schädler et al., 2019). Such an increase in temperature led to an extended frost-free period in stress-treated plots (Schädler et al., 2019). Therefore, a combination of these stress factors, which was repeated over a period of 10 years, can influence soil-associated microbial community and structure, and these effects can be amplified or mitigated depending on the type of farming management. Indeed, previous research based on the GCEF field has shown that the organic agricultural practice significantly enhanced total AMF richness under future climate (Wahdan et al., 2021). In line with this, we observed that climate history directly influenced the bacterial community structure of soil microbial extracts, which was only evident under conventional farming. For instance, the relative abundance of *Haliangiales*, *Pseudomonadales*, *Rhizobiales*, and *Sphingobacteriales* was significantly higher in ambient than in future climates. These results may indicate the increased sensitivity of some groups of bacteria to future climate changes, with more pronounced effects in conventional-based than organic farming.

In the case of treated plants, the relative abundance of *Burkholderiales* and *Sphingobacteriales* was notably higher in the future climate, but this was only observed for organic farming. Our results agree with earlier reports that *Burkholderiales* and *Sphingobacteriales* were found to be more abundant in the dry environment than in moist conditions in agricultural soil (Postma et al., 2016). Certain members of the *Burkholderiales* order, such as *Burkholderia*, can enhance plant growth and combat crop diseases through the production of various substances such as allelochemicals, antibiotics, and siderophores (Elshafie and Camele, 2021; Liu et al., 2022). Such stress-tolerant and beneficial bacteria from inoculum sourced from organic farming with the past history of future climate can potentially improve the health of indigenous seed microbiomes (and subsequently seedlings) by replacing non or less-beneficial bacterial species that are already present inside and outside of the seeds. If that is the case, such interactions between indigenous and inoculum-introduced bacteria could enhance the overall functioning of the community, leading to beneficial effects for the host plants (Afkhami, 2023; Allsup et al., 2023).

We found that seedlings exposed to soil microbes with a history of future climates in organic farming displayed a higher bacterial diversity compared to other treatments. Our results are in line with the previous findings where organic farming has been shown to promote higher soil (Lori et al., 2017; Lupatini et al., 2017; Martínez-García et al., 2018) and plant (Hartmann et al., 2015; Schmidt et al., 2019) associated microbial diversity than conventional farming. For instance, a study by Ricono *et al*. (2022) found that organic farming increased the diversity of bacterial communities in winter wheat and promoted the growth of disease-resistance bacteria (e.g., *Pseudomonadaceae*, *Burkholderiaceae*, *Xanthomonadales*, *Gammaproteobacteria*). Furthermore, our analysis of the similarity between soil microbial extracts and inoculated seedlings revealed that seedlings inoculated with water-extractable microbes from organic soil with past experience of future climate were more similar to each other than control plants. This suggests that the water-extractable microbial communities present in organic farming with a history of future climate can influence the composition of seedling bacteria. This finding is similar to previous research, indicating that soil microorganisms play an important role in the colonization of plant-associated microbiomes (Hardoim et al., 2012; Grady et al., 2019; Azarbad et al., 2020, 2022; Walsh et al., 2021). For example, Walsh et al. (2021) showed that soil microbial extracts (where soils were collected from 219 different soil types across the United States) had a significant impact on shaping the microbiomes of wheat seedlings.

Interestingly, we observed a trade-off between fast germination and seedling biomass. Seeds inoculated with microorganisms extracted from conventional agriculture under ambient climate showed the highest and fastest rate of germination. In contrast, organic agriculture with future climate-related microorganisms showed the lowest and slowest rate of germination. It is important to highlight that the combined effect of climate and agricultural management may not only affect soil biotic factors but also abiotic properties. Therefore, besides microbial effects, water-extractable nutrients from the soil, even though diluted, may also influence seedlings’ growth parameters. Indeed, we observed that conventional soil under ambient climate contained higher water extractable concentrations of total phosphorus and phosphate than other treatments. These results may, to some extent, explain the highest and fastest rate of germination in this treatment. We could not detect any pattern in terms of measured soil water extractable nutrients in organic soil under a future climate that could explain higher seedling biomass. However, we observed some interesting differences in terms of soil bacterial extracts, which may at least partly suggest the important role of microbes in enhancing seedlings above aground biomass. This suggests that conventional farming practices may prioritize rapid germination, possibly due to high inorganic fertilizer and crop protection inputs that ensure early plant establishment, particularly in ambient climates. In contrast, organic farming practices, which emphasize soil overall health, may promote the long-term development and quality of seedlings, resulting in higher above-ground biomass. However, it is important to mention that our study was performed under a controlled environment where only a proportion of soil bacteria could grow. In addition, our experiment was conducted based on one wheat genotype for a period of one week. Future studies under soil mesocosms or field conditions will help to conform our findings by taking into account more wheat genotypes and monitoring different morphological and physiological plant traits (including seed quantity and quality at harvest), together with microbial parameters throughout the growing season.

## Conclusions

Our results highlight the importance of organic farming practices in promoting bacterial diversity and potentially enhancing seedlings’ growth and biomass. This knowledge can support agricultural strategies for sustainable and resilient food production systems, especially in the face of climate uncertainty. The next step is to answer open questions regarding the details behind the functioning of the soil and plant microbiome, which are related to benefits to the plant under different histories of agricultural practices and climate.

## Acknowledgments

We would like to thank the research group at the Global Change Experimental Facility for providing us with the soil samples and access to the field. The work was conducted in Prof. Junker’s Evolutionary Ecology of Plants research group at Philipps-University Marburg.

---

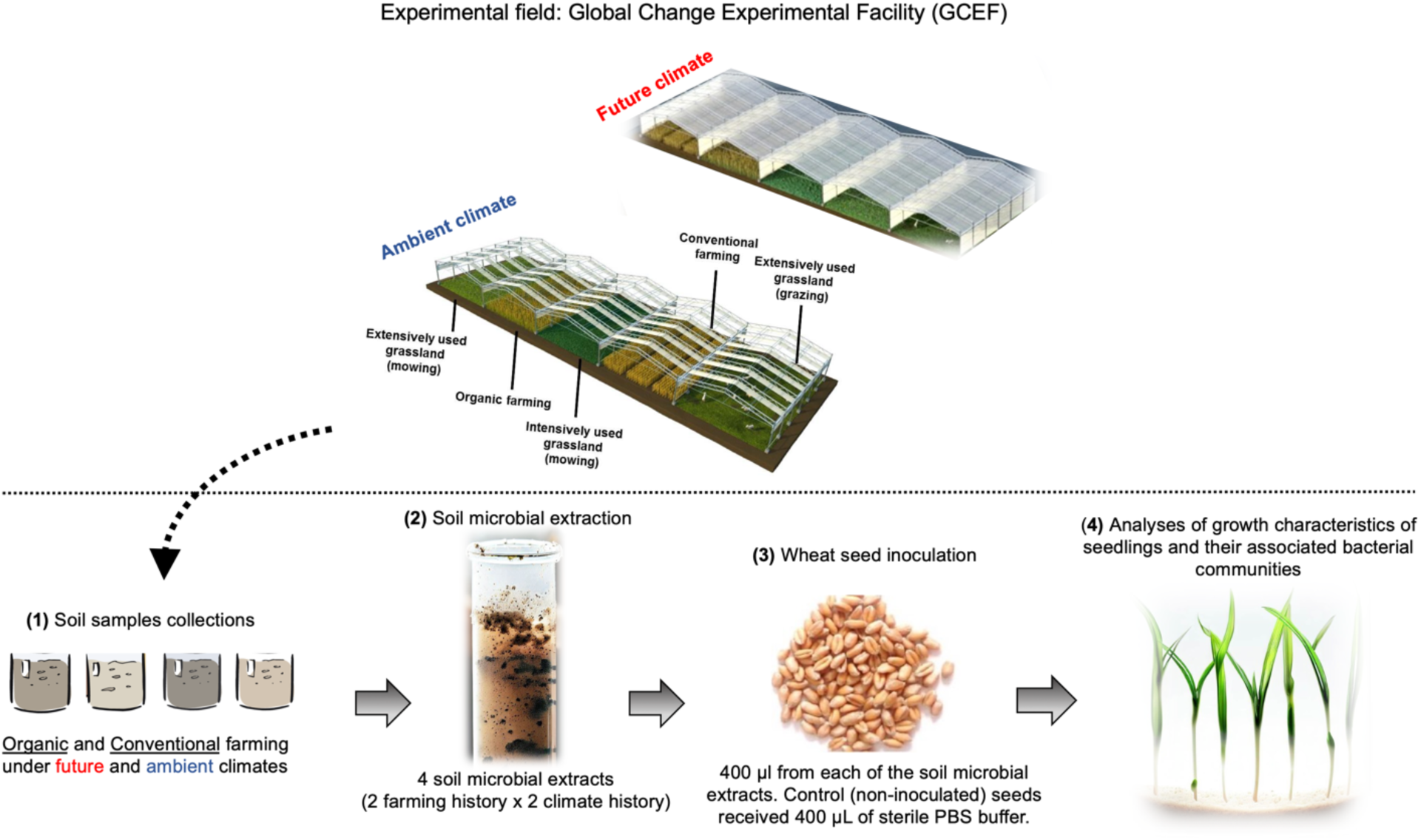
**Featured Figure.** Experimental design. (1) 20 soil sub-samples were collected from future and ambient plots associated with organic and conventional farming systems, representing different soil and microbiome stress histories. Sub-samples were mixed to have one representative sample for each farming and climate history. (2) Microbes were extracted from 5 g of each mixed soil with 45 ml of PBS buffer. (3) 6 seeds (*Triticum aestivum* L.) were placed in a sterile microbox where each seed was treated with 400 µL of soil microbial extracts on the seed. Control (non-inoculated) seeds received 400 µL of sterile PBS buffer. (4) Germination rates and germination times were measured over a period of 6 days. At the end of the experiment (8th day), the weight of the above-ground biomass of each seedling was measured for individual plants, and then shoot samples were stored at −20°C for DNA extraction and amplicon sequencing.

