## Supplementary information for "The stress history of soil bacteria under organic farming enhances the growth of wheat seedlings"

**Table S1.** Kruskal-Wallis test for the effects of farming history, climate history and their interactions on seed germination and seedlings biomass.

|  | Germination | | Biomass | |
| --- | --- | --- | --- | --- |
|  | chi-squared | p-value | chi-squared | p-value |
| Farming | 32.42 | **<0.001** | 4.371 | 0.112 |
| Climate | 35.44 | **<0.001** | 5.877 | **0.052** |
| Day | 17.9 | **0.000** | - | - |
| Farming × Climate | 72.26 | **<0.001** | 13.23 | **0.010** |
| Farming × Day | 50.90 | **<0.001** | - | - |
| Climate × Day | 53.97 | **<0.001** | - | - |
| Farming × Climate × Day | 93.80 | **<0.001** | - | - |

**Table S2.** The Bray-Curtis dissimilarity for each microbial extract and corresponding inoculated plants. The significance of the differences was tested using the Wilcoxon ranked test.

|  |  | Treated plants | | Control plants | |
| --- | --- | --- | --- | --- | --- |
| Farming history | Climate history | PC1 | PC2 | PC1 | PC2 |
| Organic | Ambient | 0.016 | 0.016 | 0.016 | 0.016 |
|  | Future | 0.036 | **0.79** | 0.036 | 0.036 |
| Conventional | Ambient | 0.015 | **0.190** | 0.015 | 0.015 |
|  | Future | 0.015 | 0.015 | 0.015 | 0.015 |

**Table S3.** ANOVA test for the effects of farming and climate histories and their interactions on the most abundant bacterial order associated with soil extracts and seedlings. Farming refers to the history of soil microbes extracted from organic and conventional agriculture, and climate refers to the history of soil microbes extracted from future and ambient climate under each farming.

.

|  | Soil extracts | | | | | | | Seedlings | | | | | |
| --- | --- | --- | --- | --- | --- | --- | --- | --- | --- | --- | --- | --- | --- |
|  | Farming | | | Climate | | Farming × Climate | | Farming | | Climate | | Farming × Climate | |
| Bacterial Order | F-value | p-value | F-value | | p-value | F-value | p-value | F-value | p-value | F-value | p-value | F-value | p-value |
| Burkholderiales | 0.435 | 0.523 | 0.985 | | 0.342 | 0.042 | 0.841 | 0.000 | 1.000 | 5.626 | **0.031** | 1.329 | 0.266 |
| Enterobacterales | 2.565 | 0.138 | 2.960 | | 0.113 | 2.537 | 0.140 | 0.016 | 0.900 | 1.639 | 0.219 | 0.961 | 0.341 |
| Flavobacteriales | 5.257 | **0.043** | 0.011 | | 0.918 | 0.210 | 0.656 | 2.472 | 0.135 | 1.369 | 0.259 | 0.139 | 0.714 |
| Gaiellales | 0.132 | 0.723 | 2.000 | | 0.185 | 0.792 | 0.392 | - | - | - | - | - | - |
| Gemmatimonadales | 0.119 | 0.737 | 0.842 | | 0.379 | 0.006 | 0.938 | - | - | - | - | - | - |
| Haliangiales | 2.586 | 0.136 | 11.403 | | **0.006** | 0.326 | 0.579 | - | - | - | - | - | - |
| Lactobacillales | - | - | - | | - | - | - | 1.040 | 0.323 | 1.015 | 0.329 | 1.047 | 0.321 |
| Micrococcales | 0.382 | 0.549 | 4.029 | | 0.070 | 0.619 | 0.448 | 2.327 | 0.147 | 1.245 | 0.281 | 1.502 | 0.238 |
| Propionibacteriales | 0.262 | 0.619 | 0.276 | | 0.610 | 1.160 | 0.305 | 1.734 | 0.206 | 0.013 | 0.911 | 0.168 | 0.687 |
| Pseudomonadales | 0.149 | 0.707 | 1.015 | | 0.335 | 3.756 | 0.079 | 2.987 | 0.103 | 0.137 | 0.716 | 0.395 | 0.538 |
| Pyrinomonadales | 0.422 | 0.529 | 0.251 | | 0.626 | 3.792 | 0.077 | - | - | - | - | - | - |
| Rhizobiales | 0.878 | 0.369 | 0.444 | | 0.519 | 0.001 | 0.972 | 0.468 | 0.504 | 0.572 | 0.460 | 0.757 | 0.397 |
| Rokubacteriales | 0.149 | 0.707 | 0.021 | | 0.886 | 0.120 | 0.735 | - | - | - | - | - | - |
| Sphingobacteriales | 0.094 | 0.765 | 1.914 | | 0.194 | 2.306 | 0.157 | 3.434 | 0.082 | 19.564 | **0.000** | 5.265 | **0.036** |
| Sphingomonadales | 0.438 | 0.522 | 0.442 | | 0.520 | 1.107 | 0.315 | 0.079 | 0.783 | 2.009 | 0.176 | 0.001 | 0.982 |
| Vicinamibacterales | 2.293 | 0.158 | 0.030 | | 0.866 | 5.807 | **0.035** | - | - | - | - | - | - |
| Xanthomonadales | 11.09 | **0.007** | 3.519 | | 0.087 | 0.687 | 0.425 | 1.166 | 0.296 | 6.058 | **0.026** | 0.000 | 0.983 |

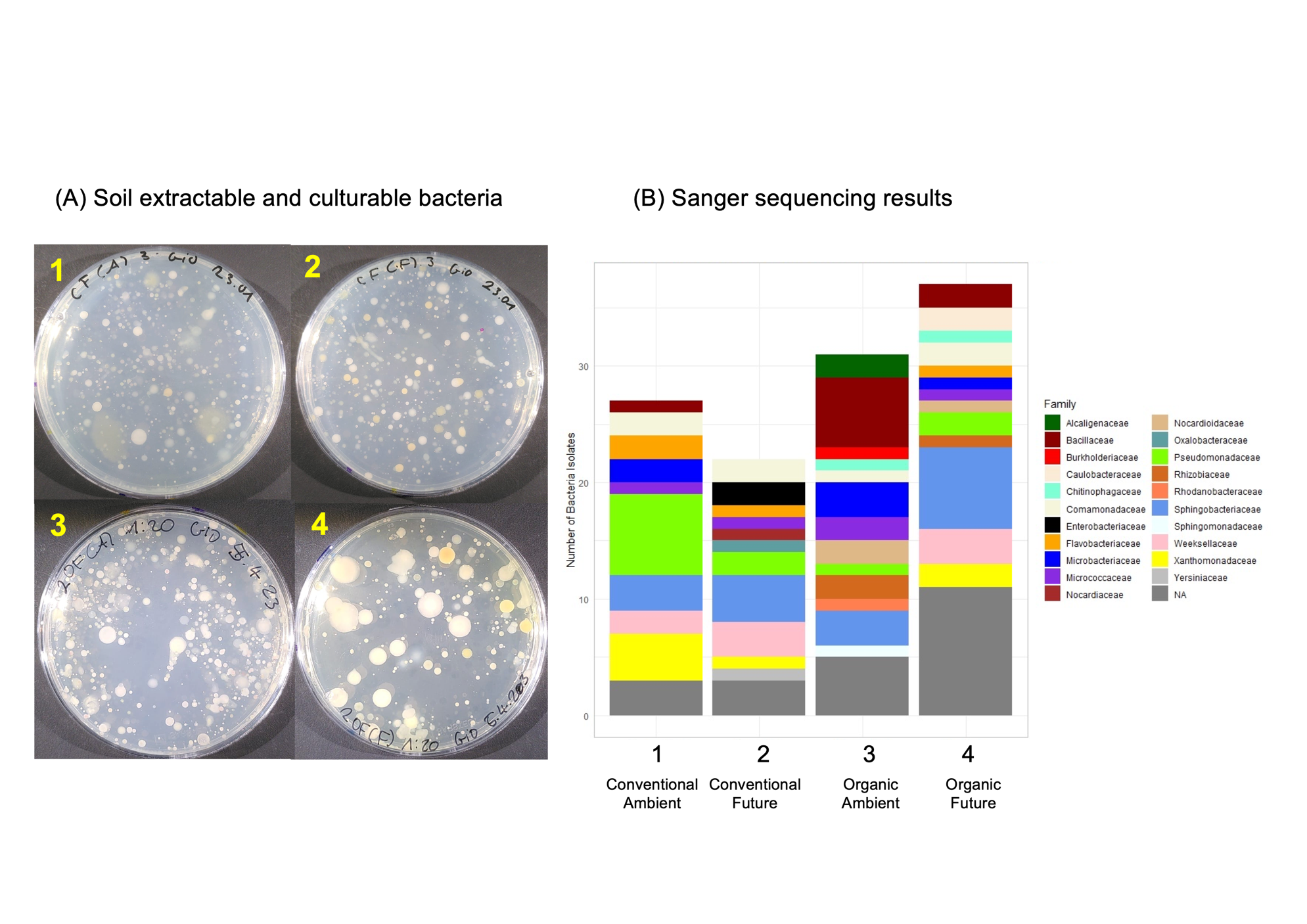

**Figure S1.** Different bacterial colonies associated with soil microbial extracts (A). The result of Sanger sequencing of the 16S of detected bacterial colonies (B).

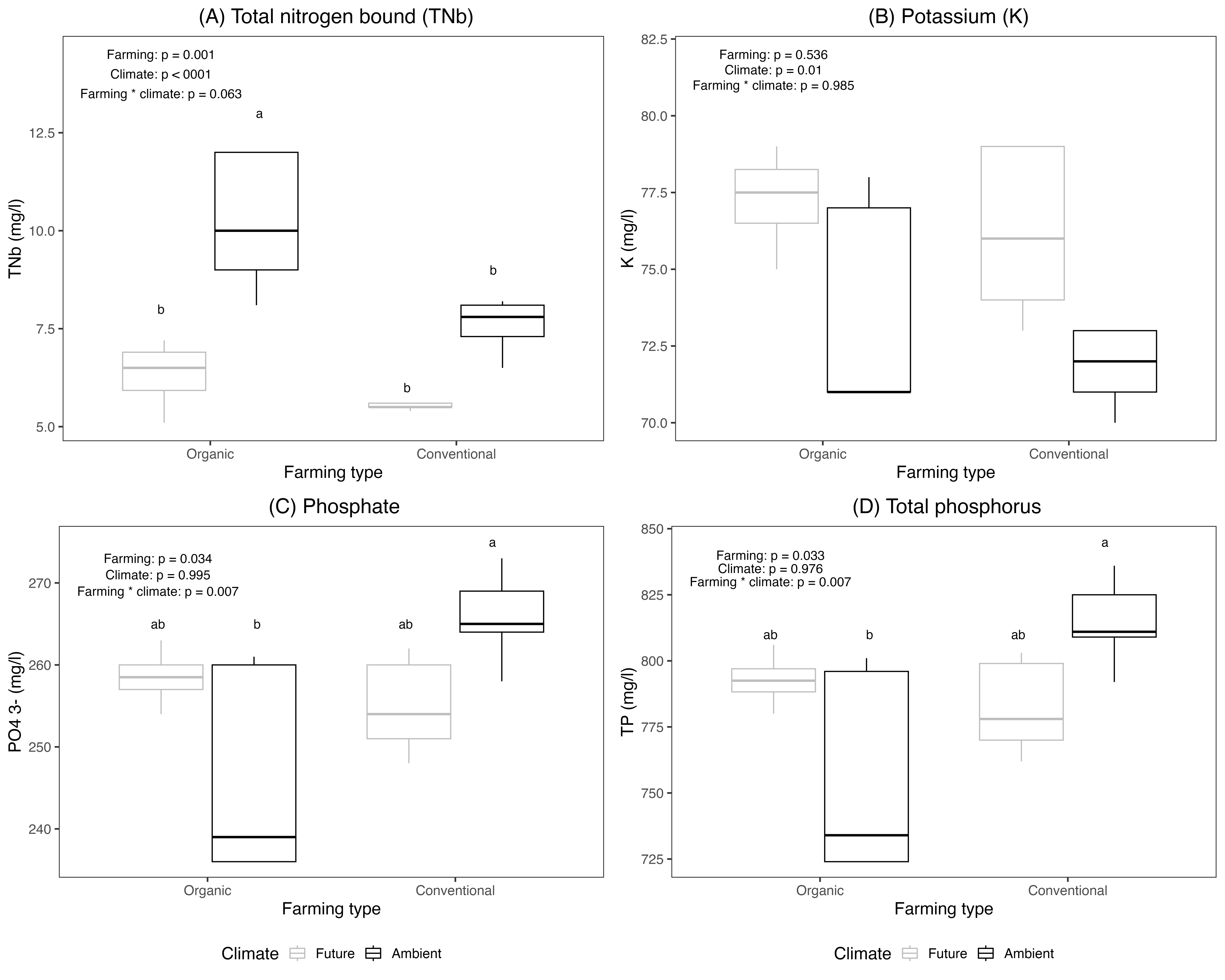

**Figure S2.** Water-extractable nutrients in soil suspensions. ANOVA was performed to test the impact of each experimental factor on water-extractable nutrients.

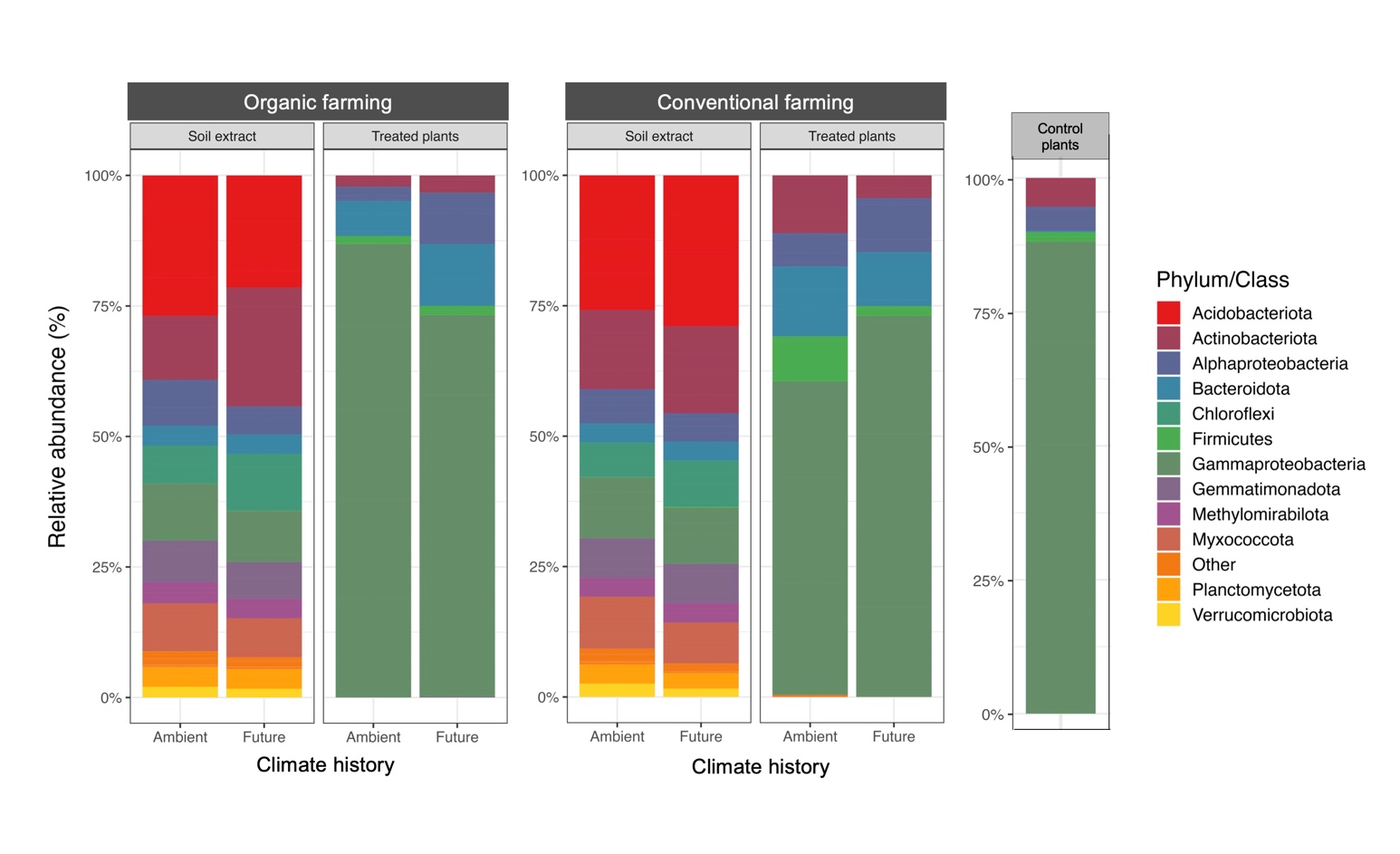

**Figure S3.** Relative abundance of the most abundant bacterial phyla (at the classes level for *Proteobacteria*) associated with soil extracts, non-inoculated (control), and inoculated (treated) plants. The relative abundance is shown according to inoculated plants and their corresponding soil microbial extracts with different farming (organic vs. conventional) and climate (future vs. ambient) histories.
